# Bidirectional transcription at the *PPP2R2B* gene locus in spinocerebellar ataxia type 12

**DOI:** 10.1101/2023.04.02.535298

**Authors:** Chengqian Zhou, Hans B. Liu, Fatemeh J. Bakhsh, Bin Wu, Mingyao Ying, Russell L. Margolis, Pan P. Li

## Abstract

**OBJECTIVE:** Spinocerebellar ataxia type 12 (SCA12) is a neurodegenerative disease caused by expansion of a CAG repeat in the *PPP2R2B gene*. Here we tested the hypothesis that the *PPP2R2B antisense* (*PPP2R2B-AS1*) transcript containing a CUG repeat is expressed and contributes to SCA12 pathogenesis.

**METHODS:** Expression of *PPP2R2B-AS1* transcript was detected in SCA12 human induced pluripotent stem cells (iPSCs), iPSC-derived NGN2 neurons, and SCA12 knock-in mouse brains using strand-specific RT-PCR (SS-RT-PCR). The tendency of expanded *PPP2R2B-AS1* (*expPPP2R2B-AS1*) RNA to form foci, a marker of toxic processes involving mutant RNAs, was examined in SCA12 cell models by fluorescence *in situ* hybridization. The toxic effect of *expPPP2R2B-AS1* transcripts on SK-N-MC neuroblastoma cells was evaluated by caspase 3/7 activity. Western blot was used to examine the expression of repeat associated non-ATG-initiated (RAN) translation of *expPPP2R2B-AS1* transcript in SK-N-MC cells.

**RESULTS:** The repeat region in *PPP2R2B* gene locus is bidirectionally transcribed in SCA12 iPSCs, iPSC-derived NGN2 neurons, and SCA12 mouse brains. Transfected *expPPP2R2B-AS1* transcripts are toxic to SK-N-MC cells, and the toxicity may be mediated, at least in part, by the RNA secondary structure. The *expPPP2R2B-AS1* transcripts form CUG RNA foci in SK-N-MC cells. *expPPP2R2B-AS1* transcript is translated in the Alanine ORF via repeat-associated non-ATG (RAN) translation, which is diminished by single nucleotide interruptions within the CUG repeat, and MBNL1 overexpression.

**INTERPRETATION:** These findings suggest that *PPP2R2B-AS1* contributes to SCA12 pathogenesis, and may therefore provide a novel therapeutic target for the disease.

## Introduction

Spinocerebellar ataxia type 12 (SCA12) is an autosomal dominant neurodegenerative disease, among the more common SCAs in India^1-3^, characterized by tremor, gait abnormalities, and neuropsychiatric syndromes^3^. Neuropathologically, the single SCA12 brain available for study revealed both cerebral cortical and cerebellar atrophy, with a noted loss of Purkinje cells^4^, in line with the cerebral cortical and cerebellar atrophy consistently observed by CT and MRI^3, 5^.

SCA12 is caused by a CAG/CTG expansion mutation in exon 7 of *PPP2R2*B, a gene encoding regulatory units of protein phosphatase 2A (PP2A)^6^. Normal alleles carry 4 to 31 triplets^7^, whereas disease alleles carry 43 to 78 triplets^2^. No polyGln aggregates were detected in the available formalin-fixed SCA12 brain, suggesting that polyGln may not contribute to SCA12 pathogenesis^4^. While all normal alleles and the majority of expanded alleles are uninterrupted, expanded alleles with single nucleotide interruptions within the CAG/CTG repeat have been reported in SCA12 patients of Uyghur ethnicity with much milder symptoms^8^, suggesting that the hairpin RNA secondary structure formed by the CAG/CUG repeat may play a contributory role in SCA12 pathogenesis. Compared with Huntington’s disease (HD) and most other CAG/CTG diseases, SCA12 has a relatively longer repeat expansion, a later disease onset, and a milder course^1^.

Natural antisense transcripts (NATs) that at least partially overlap with the sense strand gene appear to contribute to the pathogenesis of a number of diseases caused by trinucleotide repeat expansions, including fragile X syndrome (FXS) and fragile X-associated tremor/ataxia syndrome (FXTAS)^9^, Huntington’s disease (HD)^10^, Huntington’s disease-like 2 (HDL2)^11, 12^, and SCAs 2^13^, 7^14^ and 8^15^. We therefore hypothesized that in SCA12, the repeat region at *PPP2R2B* gene locus may be bidirectionally transcribed, and the antisense product may contribute to disease pathogenesis.

Here we report that the *PPP2R2B* locus is bidirectionally transcribed in SCA12 SCA12 iPSCs, iPSC derived NGN2 neurons, and SCA12 mouse brains. The antisense transcript *PPP2R2B-AS1* with a CUG repeat expansion is neurotoxic, and therefore may contribute to SCA12 pathology. This finding suggests that *PPP2R2B-AS1* is a potential therapeutic target in SCA12.

## Results

### Bidirectional transcription at the PPP2R2B gene locus

Modeled on our previous work on HDL2 and SCA2^11-13^, diseases in which a relatively short repeat expansion triggers a devastating phenotype, probably through a combination of loss-of-function, RNA-mediated, and protein-mediated toxicity, we speculated that SCA12 may similarly involve complex mechanisms of pathogenesis. We therefore tested the possibility that an antisense transcript with an expanded CUG repeat is expressed from the opposite strand of *PPP2R2B*. In conjunction with Cedars Sinai Stem Cell Core, we generated and characterized 8 iPSC lines derived from the fibroblasts of three different human SCA12 patients (Table 1), while three matched control iPSC lines were included for this study. We differentiated control and SCA12 iPSCs into cortical excitatory neurons using an NGN2 overexpression protocol^16, 17^. Using total RNA extracted from control and SCA12 iPSCs as well as iPSC-derived NGN2 neurons, we performed strand-specific reverse transcription PCR (SS-RT-PCR)^13^ with linkered (LK) primers flanking the repeat in *PPP2R2B* exon 7 (Fig. 1A). We detected the expression of *PPP2R2B* antisense transcripts from both normal and expanded alleles from SCA12 iPSCs or iPSC derived NGN2 neurons, and from the normal alleles in the corresponding controls (Fig. 1B and 1C). In keeping with HUGO guidelines, we named the gene expressing this transcript *PPP2R2B* antisense 1, *PPP2R2B-AS1*. The identity of the transcript was confirmed by sequencing (data not included). Next, we examined the expression of *PPP2R2B-AS1* in a novel humanized SCA12 knock-in mouse model that was generated by replacing the mouse *PPP2R2B* exon 2 containing a short, interrupted repeat with its human counterpart, exon 7, with either 10 or 80 CAG triplets (KI-10 or KI-80; Li and Margolis, unpublished results). SS-RT-PCR detected expanded *PPP2R2B-AS1* (*expPPP2R2B-AS1*) transcripts in the cortices and cerebella of three-month-old male homozygous SCA12 KI-80 mice, while as expected, normal *PPP2R2B-AS1* (*nPPP2R2B-AS1*) transcripts were expressed in the age- and sex-matched homozygous SCA12 KI-10 or wildtype (WT) cortices or cerebella (Fig. 1D). The *PPP2R2B-AS1* transcript fragment detected by SS-RT-PCR contains 44bp upstream and 19bp downstream of the CUG repeat, and includes no ATG start codons.

**Table 1.**
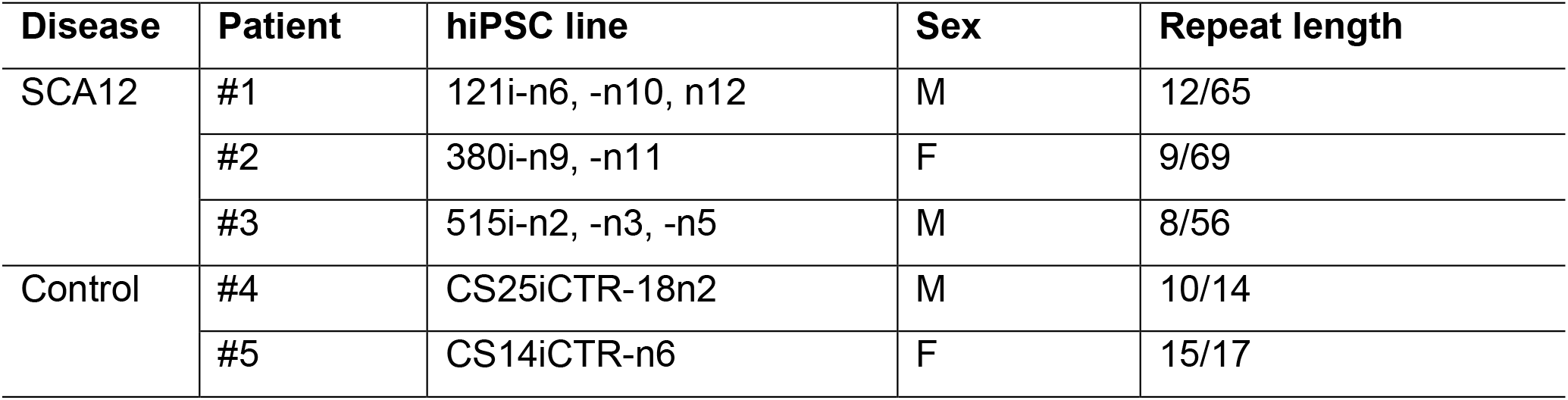
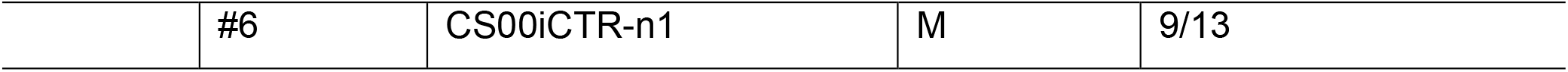
Human SCA12 and control iPSC lines used.

**Fig. 1.**
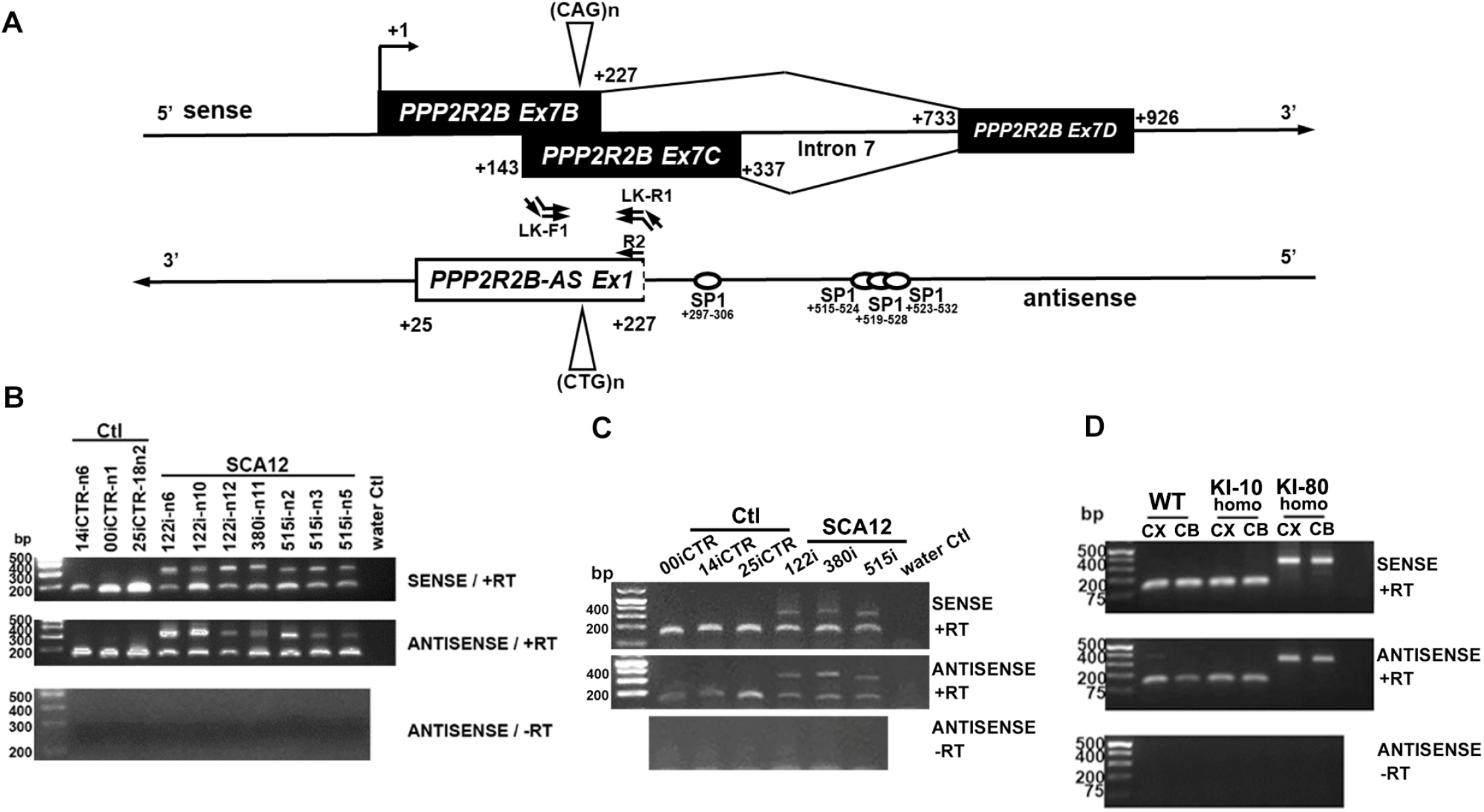
Bidirectional transcription at *PPP2R2B* gene locus. (A) *PPP2R2B-AS1* map. Primer locations for SS-RT-PCR in B-D are shown. On the sense strand, *7B7D* and *7C7D* are the two splice variants of *PPP2R2B* exon 7 that contains the CAG repeat. On the antisense strand, location of SP1 binding sites were predicted using PROMO^18^. 3’ end of *PPP2R2B-AS1* was identified by 3’ RACE. The 5’ end of *PPP2R2B-AS1* exon 1 is not characterized. (B-D) Detection of normal and expanded *PPP2R2B* and *PPP2R2B-AS1* by SS-RT-PCR in human iPSCs (B), iPSC derived cortical excitatory neurons differentiated by NGN2 overexpression (C), and cortices (CX) and cerebella (CB) of a humanized SCA12 knock-in mouse model with either 10 or 80 repeats (KI-10 or KI-80). PCR bands amplified from two normal alleles in human control iPSCs or iPSC derived neurons could not be separated on an agarose gel due to size similarity. Only homozygous (homo) mice were included. N=3 independent experiments; representative gel images are shown.

3’ RACE was performed to characterize the 3’ end of the *PPP2R2B-AS1* transcript in control human brain, and we detected additional 3’ sequence (145bp downstream of the CTG repeat; without ATGs) followed by a polyA tail, indicating that the *PPP2R2B-AS1* transcript is indeed polyadenylated (Fig. 1A). Despite multiple attempts at 5’ RACE, the 5’ end remains uncharacterized, presumably because high GC content prevented successful PCR amplification. Nevertheless, three SP1 binding sites are predicted in the region 98∼324 bp upstream of the CTG repeat^18^, indicating potential transcription initiation sites for the *PPP2R2B-AS1* transcript. Future full-length RNAseq using control human brain tissue, or SCA12 KI mouse brain, may reveal the entire sequence and possible splicing isoforms of *PPP2R2B-AS1*, and guide experiments to confirm the presence and properties of a functional promoter for *PPP2R2B-AS1*, and suggest likely ORFs (if any) within the full length *PPP2R2B-AS1* transcript.

### expPPP2R2B-AS1 triggers toxicity in neuroblastoma SK-N-MC cells

Transcripts containing expanded CUG repeats contribute to toxicity in DM1, HDL2, and SCA2^11, 13^, and perhaps other CAG/CTG diseases^19^. We therefore examined whether *expPPP2R2B-AS1* with a CUG repeat expansion within the physiological range of adult-onset SCA12 is toxic to neuronal-like cells. *PPP2R2B-AS1-(CTG)n* constructs expressing *PPP2R2B-AS1* transcript fragments with either 10 or 73 CUG triplets and flanking regions extending 44 bp upstream and 145 bp downstream from the repeat were transfected into SK-N-MC neuroblastoma cells. The toxicity of exogenous *PPP2R2B-AS1* was compared to that of the exogenous *ATXN2 antisense* (*ATXN2-AS)* transcript fragment with 44 (vs. 22) CUG triplets, previously shown to be associated with SCA2 pathogenesis and toxic to SK-N-MC cells^13^. *PPP2R2B-AS1-(CTG)73* triggered significant toxicity that was similar to that of the *ATXN2-AS-(CTG)44* transcript and was about 1.5 times more toxic than *PPP2R2B-AS1-(CTG)10 and ATXN2-AS-(CTG)22* (Fig. 2A).

**Fig. 2.**
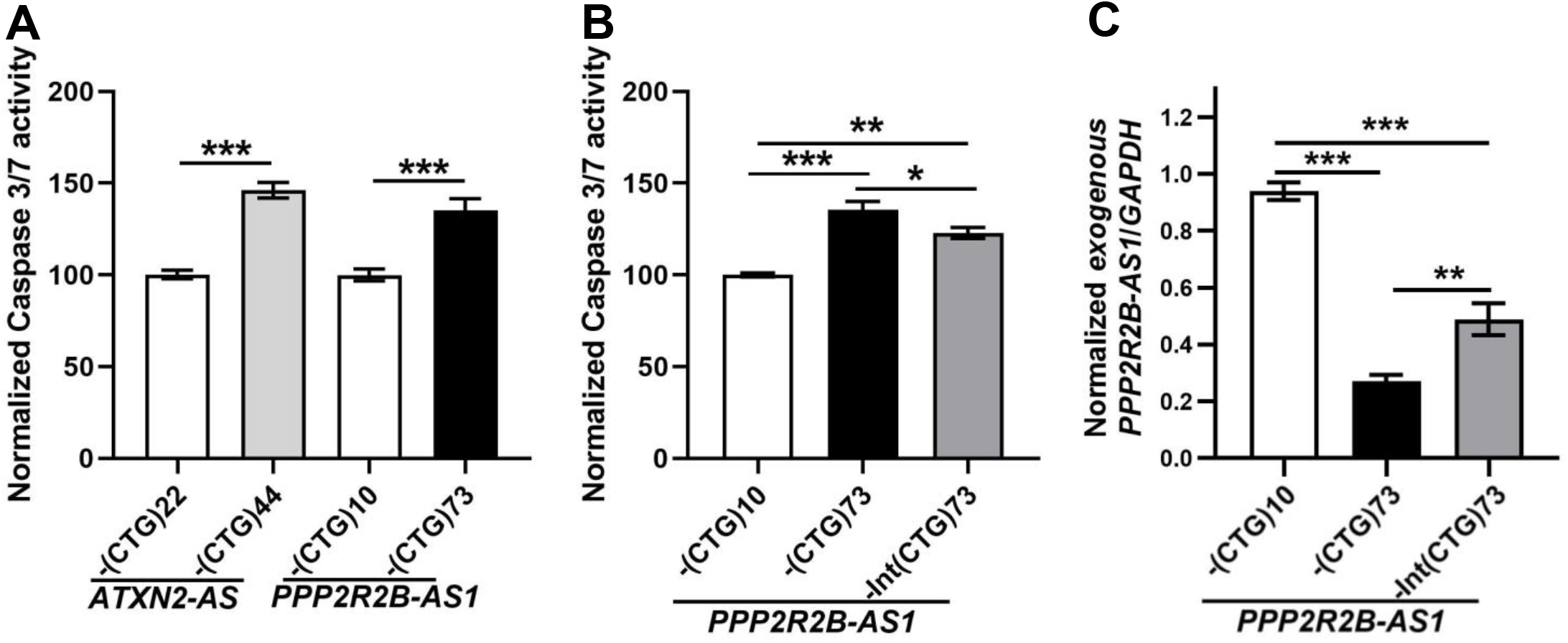
Overexpression of an expanded *PPP2R2B-AS1* transcript is toxic to SK-N-MC cells. (A) Caspase 3/7 assay indicates toxicity of an expanded *PPP2R2B-AS1-(CUG)73* transcript compared to a normal *PPP2R2B-AS1-(CUG)10* transcript. *ATXN2-AS* constructs are included as positive controls. N=5 biological replicates. The Caspase 3/7 activity in *ATXN2-AS-(CTG)22* or *PPP2R2B-AS1-(CTG)10* transfected SK-N-MC cells was normalized to 100. (B) Single nucleotide interruptions in the CUG repeat decrease the toxicity of expanded *PPP2R2B-AS1* transcript. N=4 biological replicates. The Caspase 3/7 activity in *PPP2R2B-AS1-(CTG)10* transfected SK-N-MC cells was normalized to 100. (C) Relative expression level of exogenous *PPP2R2B-AS1* transcripts in SK-N-MC cells normalized by *GADPH* level. qPCR primers used to measure exogenous *PPP2R2B-AS1* are located on the vector backbone. N=3 biological replicates. The *PPP2R2B-AS1*/*GAPDH* in *PPP2R2B-AS1-(CTG)10* transfected SK-N-MC cells was normalized to 1. Mean ± SEM are shown. One-way ANOVA and Tukey posthoc test, *p<0.05, **p<0.01, ***p<0.001.

Individuals from a single SCA12 pedigree of Uyghur ethnicity are predicted to have an expanded SCA12 allele that would express *expPPP2R2B-AS1* with 1-3 single nucleotide interruptions in the repeat^8^; these individuals have a mild form of SCA12^8^, suggesting that interruptions in the CUG repeat of *expPPP2R2B-AS*1 may reduce neurotoxicity. It has been suggested that CUG transcript toxicity is dependent on the hairpin structure formed by the repeat^13^, the stem of which sequesters RNA-binding proteins (RBPs), i.e. MBNL1; we therefore hypothesized that the *expPPP2R2B-AS1* transcripts with an interrupted repeat, as found in the Uyghur family, may be less toxic than *expPPP2R2B-AS1* transcripts with an uninterrupted CUG repeat of the same length. To test this hypothesis, we introduced three single nucleotide mutations into the repeat of *PPP2R2R-AS1-(CTG)73*, in order to obtain *PPP2R2B-AS1-Int(CTG)73*, with a repeat composition of CTG)_23_CTC(CTG)_14_CCG(CTG)_13_TTG(CTG)_20_, similar to the interrupted *expPPP2R2B-AS1* transcript predicted in the Uyghur population. Inserting interruptions indeed decreased *expPPP2R2B-AS1* toxicity in SK-N-MC cells (Fig. 2B). The levels of overexpressed transcripts in SK-N-MC cells were measured by quantitative RT-PCR (qRT-PCR; Fig. 2C), indicating that though the same amount (1ug) of dsDNA plasmid of each construct was transfected, the *PPP2R2B-AS1-(CUG)73* transcript was expressed at a much lower level than *PPP2R2B-AS1-Int(CUG)73* and *PPP2R2B-AS1-(CUG)10*, suggesting that the *expPPP2R2B-AS1* toxicity observed in the Caspase 3/7 assay may be underestimated. We speculate that the low level of *PPP2R2B-AS1-(CUG)73* expression may be the consequence of nuclear retention, perhaps in foci (see below). This set of experiments suggests that the toxicity of *expPPP2R2B-AS1* is dependent on both repeat length and repeat composition, potentially related to the secondary structure formed by the repeat structures.

### expPPP2R2B-AS1 transcripts form CUG RNA foci

Repeat-containing mutant transcripts form RNA foci in all CUG/CAG diseases in which RNA neurotoxicity contributes to pathogenesis ^20-22^. RNA foci are also a hallmark of RNA toxicity in neurodegenerative repeat diseases associated with other types of repeats, including the ATTCT pentamer expansion that causes spinocerebellar ataxia type 10 (SCA10) ^23^ and the GGGGCC hexamer expansion in c9orf72 that is associated with ALS ^24^.

We therefore sought to detect similar foci in *PPP2R2B-AS1* overexpressing SK-N-MC cells by fluorescence *in situ* hybridization (FISH) using a 20-mer CAG LNA/DNA probe that binds to the CUG repeat^25^. CUG RNA foci were absent in SK-N-MC neuroblastoma cells that overexpress *PPP2R2B-AS1-(CTG)10* (Fig. 3A), but were not uncommon in cells overexpressing *PPP2R2B-AS1-(CTG)73* (Fig. 3B) or *PPP2R2B-AS1-Int(CTG)73* (Fig. 3C), as quantified in Fig. 3M. *PPP2R2B-AS1-(CUG)73* and *PPP2R2B-AS1-Int(CUG)73* formed similar number of foci, except that the % cells with 10+ foci was higher in *PPP2R2B-AS1-(CUG)73* cells, compared with *PPP2R2B-AS1-Int(CUG)73* cells. This set of experiments demonstrates that *expPPP2R2B-AS1* transcripts form RNA foci.

**Fig. 3.**
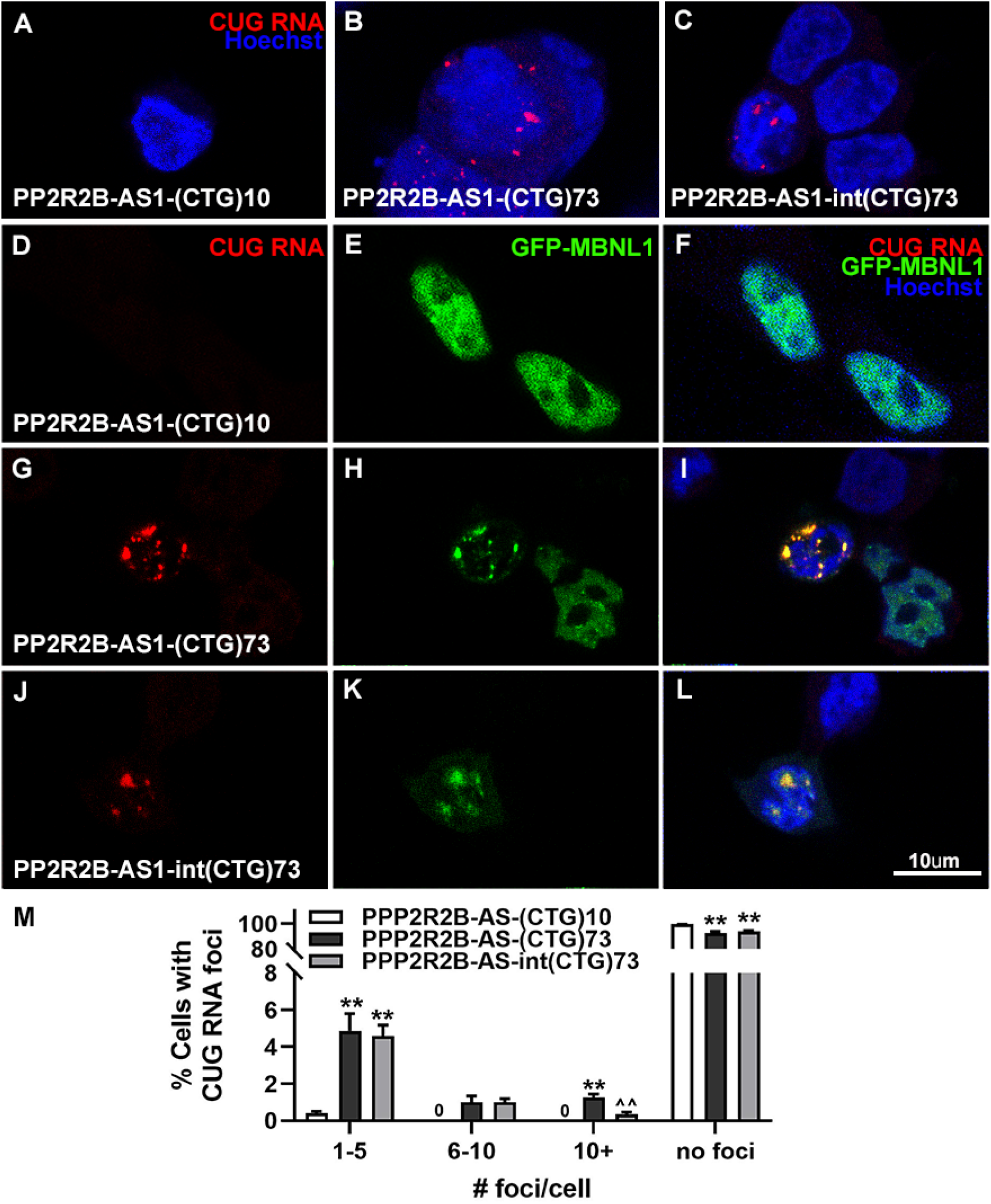
Expanded *PPP2R2B-AS1* transcripts form CUG RNA foci. (A-C) In situ hybridization using a CAG DNA/LNA probe indicates *PPP2R2R-AS1-(CUG)73* and *–int(CUG)73* transcripts form nuclear CUG RNA foci when transfected into SK-N-MC neuroblastoma cells. % of cells with foci and number of foci per cell were quantified (M). Mean ± SEM are shown. N=3 biological replicates, One-way ANOVA, *p<0.05, **p<0.01, compared with *PPP2R2B-AS1-(CTG)10*, ^^p<0.01, compared with *PPP2R2B-AS1-(CTG)73*. (D-L) GFP-MBNL1 shows nuclear diffused localization in cells that overexpress *PPP2R2B-AS1-(CUG)10* (D-F), but becomes sequestered to CUG RNA foci formed by *PPP2R2B-AS1-(CUG)73* (G-I) or *-Int(CUG)73* (J-L) transcripts. Scale bar = 10um.

Consistent with previous reports that MBNL1, a nuclear splicing factor that aberrantly interacts with CUG repeat transcripts, colocalizes to CUG RNA foci^13, 26, 27^, we also found that although exogenously expressed GFP-MBNL1 is diffusely localized to the nuclei of SK-N-MC cells co-transfected with *PPP2R2B-AS1-(CTG)10* (Fig. 3F), co-expression of GFP-MBNL1 and *PPP2R2B-AS1-(CTG)73 or PPP2R2B-AS1-Int(CTG)73* results in co-localization of MBNL1 with CUG RNA foci (Fig. 3I and 3L). While the toxicity of RNA foci themselves remains to be determined, the colocalization of RNA foci with MBNL1 in this model is suggestive of a pathogenic role of *expPPP2R2B-AS1* in SCA12.

### RAN translation of the expPPP2R2B-AS1 transcript

The experimentally-defined transcript sequence of *PPP2R2R-AS1* (the sequence used in our experiments) does not include ATG start codons upstream of the CUG repeat and would not be predicted to yield protein products via canonical ATG-initiated translation. However, hairpin-forming expanded CUG repeats can also be translated in the absence of ATG start codon through repeat-associated non-ATG translation (RAN translation) ^28^. To determine if *PPP2R2B-AS1* has the potential of undergoing RAN translation, and if RAN-translated protein fragments with expanded amino acid tracts lead to neurotoxicity, we cloned an *PPP2R2B-AS1* fragment containing multiple upstream stop codons, and a CTG repeat expansion with flanking regions extending 44 bp upstream and 26 bp downstream from the repeat into a vector with tags for each of the three open reading frames (Fig. 4A), to obtain the *PPP2R2B-AS1-(CTG)n-3T* and *PPP2R2B-AS1-Int(CTG)73-3T* constructs. Immunostaining and western blot using antibodies against each epitope tag detected a polyalanine-containing RAN product from SK-N-MC cells that overexpress *PPP2R2B-AS1-(CTG)73-3T* (Fig. 4C and 4J) or *PPP2R2B-AS1-Int(CTG)73-3T* (Fig. 4D and 4J), but not polyleucine-(Fig. 4B-D) or polycystine-containing (Fig. 4F-H) RAN products. Quantification of polyalanine RAN protein levels reveals that the *PPP2R2B-AS-Int(CTG)73-3T* construct produced less polyalanine RAN protein than the *PPP2R2B-AS1-(CTG)73-3T* construct (Fig. 4K), indicating that the expression level of polyalanine RAN product is likely dependent on the secondary structure of the *expPPP2R2B-AS1* transcript.

**Fig. 4.**
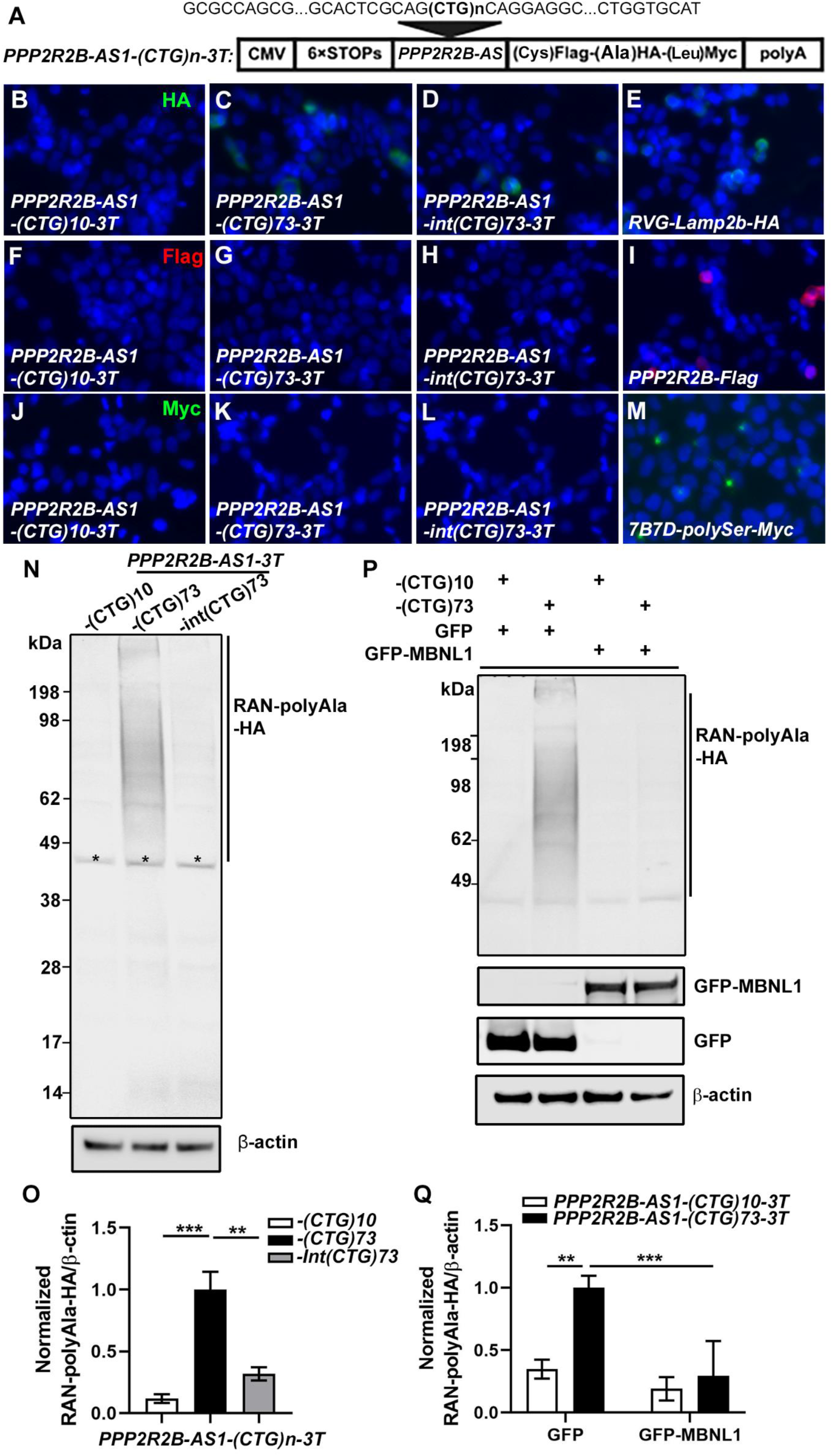
The expanded CUG repeat in *PPP2R2B-AS1* transcript may be translated into polyalanine (polyAla) containing proteins via RAN translation. (A) Schematic presentation of the tagged *PPP2RAB-AS*-(CTG)n-3T construct to determine the presence of RAN product. (B-I) Anti-HA staining revealed polyAla RAN protein in *PPP2R2B-AS1-(CTG)73-3T* and *PPP2R2B-AS1-int(CTG)73-3T* overexpressing SK-N-MC cells (Fig. 4C and 4D), compared with the *PPP2R2B-AS1-(CTG)10-3T* control cells (Fig. 4B), while anti-Flag or anti-Myc staining showed negative results (Fig. 4F-4H and 4J-4L). SK-N-MC cells transfected with *RVG-Lamp2b-HA* (Fig. 4E), *PPP2R2B-Flag* (Fig. 4I), and *7B7D-polySer-Myc* (Fig. 4M), were used as positive controls for immunostaining using anti-HA, Flag, and Myc tag antibodies, respectively. (J-K) RAN-polyAla-HA protein expression is decreased in *PPP2R2B-AS1-int(CTG)73-3T* overexpressing cells, compared with *PPP2R2B-AS1-(CTG)73-3T*. Asterisks mark nonspecific bands. N=4 biological replicates. The RAN-polyAla-HA/β-actin ratio in *PPP2R2B-AS1-(CTG)73-3T* transfected SK-N-MC cells was normalized to 1. Mean ± SEM are shown. (L-M) GFP-MBNL1 overexpression decreased RAN-polyAla-HA expression in *PPP2R2B-AS1-(CTG)73* expressing SK-N-MC cells. N=3 biological replicates. The RAN-polyAla-HA/β-actin ratio in SK-N-MC cells co-transfected with *PPP2R2B-AS1-(CTG)73-3T* and *GFP* was normalized to 1. Mean ± SEM are shown. One-way ANOVA and Tukey posthoc test, *p<0.05, **p<0.01, ***p<0.001.

### *MBNL1 decreases RAN translation of the* expPPP2R2B-AS1 *transcript*

Given our findings that GFP-MBNL1 is sequestered to the CUG RNA foci formed by the *expPPP2R2B-AS1* transcripts (Fig. 3G-3L), and previous reports that overexpression of MBNL1 decreased protein translation of transcripts containing expanded *CAG/CUG* (*expCAG/CUG)* repeat^29^, we therefore hypothesized that the MBNL1, via its interaction with the expanded *CUG* repeat, may inhibit the translation of polyalanine RAN protein from the *expPPP2R2B-AS1* transcript. In support of our hypothesis, we found that overexpression of GFP-MBNL1 (vs. GFP alone) decreased the expression of polyalanine RAN protein from the *PPP2R2B-AS1-(CUG)73-3T* transcript in SK-N-MC cells (Fig. 4L and M). The MBNL1 protein consists of four zinc finger (ZNF) domains, an unstructured domain, and a C-terminal splicing domain containing the nuclear localization signal (NLS)^30, 31^ (Fig. 5A). It was previously reported that the CUG repeats interact with MBNL1 via its N-terminal ZNF domains. To ascertain whether the effect of MBNL1 on polyalanine RAN protein production is dependent on the ZNF domains, we obtained constructs encoding GFP-tagged truncated MBNL1 proteins lacking either the C-terminal splicing domain (GFP-MBNL1-Δ251-388), or all four ZNF domains in the N-terminus (GFP-MBNL1-Δ1-240) (Fig. 5A). GFP-MBNL1 (Fig. 5B and 5E) and GFP-MBNL1-Δ1-240 (Fig. 5J and 5M) are both diffusedly localized in the nucleus, while GFP-MBNL1-Δ251-388, due to the lack of C-terminal domain containing the NLS, is expressed diffusely in nucleus and cytoplasm (Fig. 5F and 5I). GFP tagged full-length or truncated MBNL1 and *PPP2R2B-AS1* were overexpressed in SK-N-MC cells, and FISH experiments revealed that similar to full length GFP-MBNL1 (Fig. 5C-E), GFP-MBNL1-Δ251-388 (truncated MBNL1 containing ZNFs) is still colocalized with CUG RNA foci formed by *PPP2R2B-AS1-(CUG)73* transcripts (Fig. 5G-I), while GFP-MBNL1-Δ1-240 was detected diffusely in the nucleus with no colocalization with CUG RNA foci (Fig. 5K-M), confirming that the ZNF domains of MBNL1 are responsible for interacting with transcripts containing expanded CUG repeats. Next, we examined the effect of truncated MBNL1 on the production of polyalanine RAN protein from the *expPPP2R2B-AS1-3T* transcripts. While overexpression of GFP-MBNL1-Δ251-388 decreased the production of the polyalanine RAN protein from *PPP2R2B-AS1-(CUG)73-3T* transcripts, mimicking the effect of full length GFP-MBNL1, overexpression of GFP-MBNL1-Δ1-240 (lacking all the four ZNF domains) did not affect polyalanine RAN protein production (Fig. 5N and 5O), indicating that ZNF domains are necessary for MBNL1 to inhibit RAN translation of the *expPPP2R2B-AS1* transcript. However, not all four ZNF domains may be necessary, as overexpression of GFP-MBNL1-Δ12-46, a GFP tagged MBNL1 lacking the first ZNF only (Fig. S1A), also decreased the RAN polyalanine production from the *PPP2R2B-AS1-(CUG)73-3T* transcripts (Fig. S1B and S1C), mimicking the effect of full length GFP-MBNL1. To ascertain whether the inhibitory effect of MBNL1 ZNF domains on translation of the *expanded CUG repeat* transcripts also applies to ATG-initiated translation, we included a construct of *JPH3* exon 2A containing an expanded CTG repeat (relevant to HDL2), *JPH3 Ex2A-(CTG)55*, which encodes a protein containing a long polyalanine tract via ATG translation^11, 29^. Similar to the RAN translation of the *PPP2R2B-AS1-(CUG)73-3T* transcripts, overexpression of GFP-MBNL1 (Fig. 5P), GFP-MBNL1-Δ251-388 (Fig. 5P), GFP-MBNL1-Δ12-46 (Fig. S1B), but not that of GFP-MBNL1-Δ1-240 (Fig. 5P) decreased the expression of polyalanine protein from the *JPH3 Ex2A-(CTG)55* (Fig. 5Q and Fig. S1C), indicating that MBNL1 ZNF domains have inhibitory effects on the ATG-translation of the expanded *CUG* repeat as well.

**Fig. 5.**
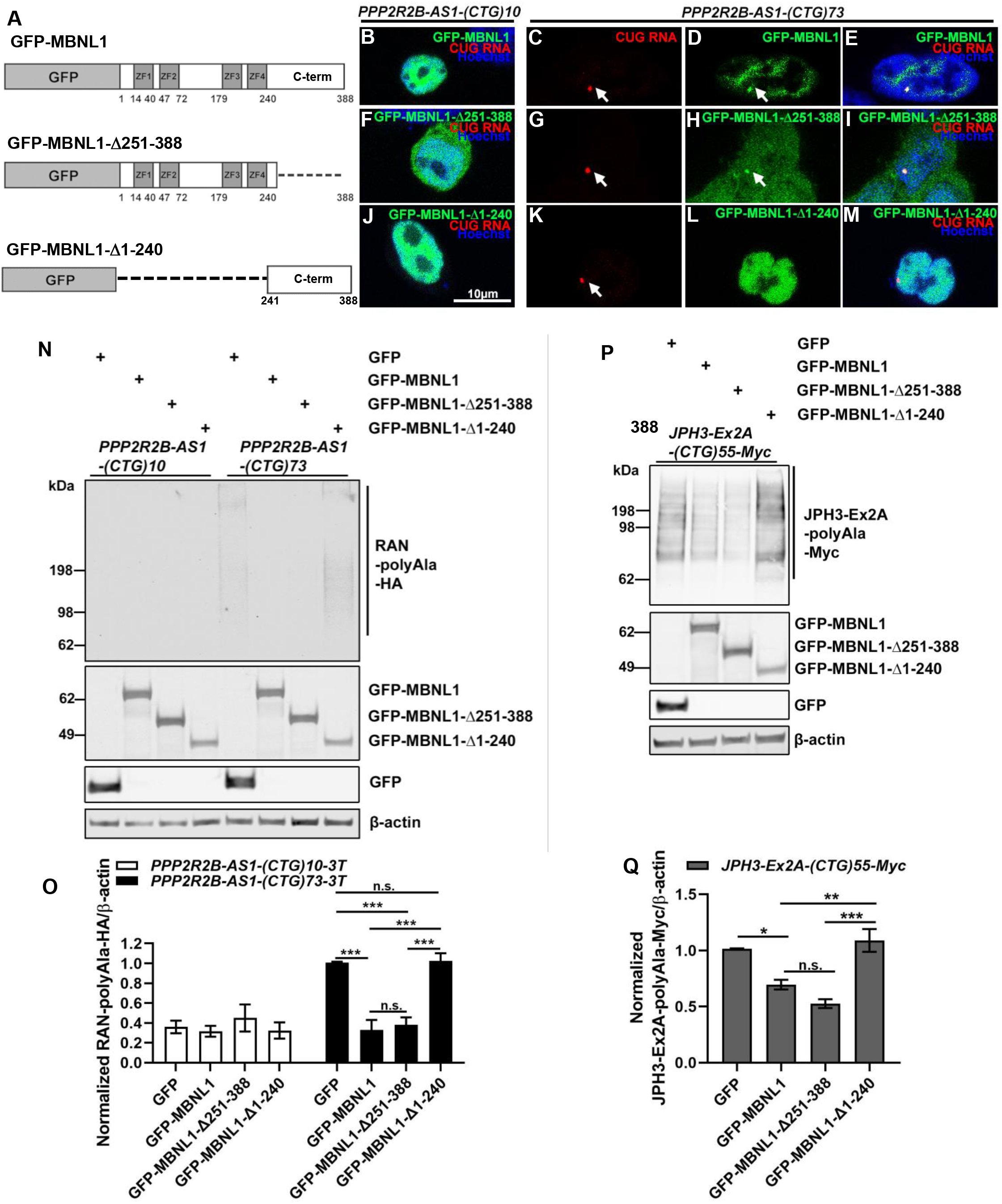
Effect of MBNL1 ZNFs on the production of RAN polyAla proteins from the *expPPP2R2B-AS1* transcript. (A). Schematic presentation of the GFP-MBNL1, GFP-MBNL1-Δ251-388, and GFP-MBNL1-Δ1-240 constructs. (B-M). *In situ* hybridization using a CAG DNA/LNA probe indicates exogenous GFP-MBNL1 and GFP-MBNL1-Δ251-388, but not GFP-MBNL1-Δ1-240, colocalizes with nuclear CUG RNA foci formed by the *PPP2R2R-AS1-(CUG)73* transcripts in SK-N-MC neuroblastoma cells. N=3 biological replicates. Representative images are shown. Scale bar = 10um. Arrows point to CUG RNA foci. (N-O) Overexpression of GFP-MBNL1 or GFP-MBNL1-Δ251-388, but not that of GFP-MBNL1-Δ1-240, decreased RAN-polyAla-HA protein expression in *PPP2R2B-AS1-(CTG)73-3T* expressing SK-N-MC cells. N=4 biological replicates. The RAN-polyAla-HA/β-actin ratio in SK-N-MC cells co-transfected with *PPP2R2B-AS1-(CTG)73-3T* and *GFP* was normalized to 1. Mean ± SEM are shown. One-way ANOVA and Tukey posthoc test, *p<0.05, **p<0.01, ***p<0.001. (P-Q) Overexpression of GFP-MBNL1, or GFP-MBNL1-Δ251-388, but not that of GFP-MBNL1-Δ1-240, decreased polyAla-Myc expression in *JPH3-Ex2A-(CTG)55-Myc* expressing SK-N-MC cells. N=4 biological replicates, Mean ± SEM are shown. The JPH3-Ex2A-polyAla-Myc/β-actin ratio in SK-N-MC cells co-transfected with *JPH3-Ex2A-(CTG)55-Myc* and *GFP* was normalized to 1. One-way ANOVA and Tukey posthoc test, *p<0.05, **p<0.01, ***p<0.001.

## Discussion

We demonstrate that both normal and expanded CUG repeat-containing *PPP2R2B-AS1* transcripts are expressed in human SCA12 iPSCs and iPSC derived cortical neurons, as well as in a novel SCA12 knock-in mouse model. The *expPPP2R2B-AS1* transcripts form foci that sequester the splicing factor MBNL1. Further, we show that the *expPPP2R2B-AS1* with a CUG repeat size within the physiological range for adult-onset SCA12 is toxic to neuronal-like SK-N-MC neuroblastoma cells. Finally, the *expPPP2R2B-AS1* transcript expresses polyalanine RAN protein which is decreased by single nucleotide interruptions within the CUG repeat, overexpression of full-length MBNL1, or overexpression of a truncated MBNL1 containing ZNF domains. We conclude that the *expPPP2R2B-AS1* transcripts may play a role in the pathogenesis of SCA12.

In DM1, and HD, normal, but not expanded antisense transcripts are expressed at the respective disease loci in human tissue^10, 11, 32^. In HDL2, expanded antisense transcript was detected in a BAC-HDL2 transgenic mouse model^12^, but not in frozen human postmortem brains^11^. In SCA2 and SCA8, the antisense transcript with the expanded repeat is expressed at significant levels^13, 15^. Only one human SCA12 postmortem brain (formalin fixed) was available for this study^33^, and SS-RT-PCR detected only the antisense allele with the normal repeat (data not included). We speculate that detection of the low-abundant antisense transcript with the expanded repeat likely requires better quality RNA than is available from the formalin fixed brain; we anticipate the availability of such tissue in the future. Nevertheless, *expPPP2R2B-AS1* was detected in undifferentiated and differentiated SCA12 iPSCs (Fig. 1B and 1C), and in the cortices and cerebella of SCA12 knock-in mice (Fig. 1D), indicating the potential relevance of *expPPP2R2B-AS1* to SCA12.

NATs frequently regulate sense strand transcript expression^37,23^, as in the case of *HTT-AS* regulation of *HTT* ^10, 34^. Our preliminary data suggests that overexpression of a *PPP2R2B-AS1* fragment *in trans* does not have a substantial effect on the expression of endogenous *PPP2R2B* transcript in SK-N-MC neuroblastoma cells (data not shown). Whether *PPP2R2B-AS1* affects the expression of *PPP2R2B in cis* remains to be determined.

Overexpressed *expPPP2R2B-AS1* fragments readily aggregate into RNA foci in SK-N-MC cells (Fig. 3). The role of these foci in the pathogenesis of other repeat expansion diseases remains controversial^13, 35, 36^; it is not fully established whether the foci themselves are protective, an epiphenomenon or neurotoxic. Similarly, the direct effect of the foci in SCA12 pathogenesis awaits further elucidation.

Our data nevertheless indicate a toxic role of *expPPP2R2B-AS1* transcripts. Although the mechanism of *expPPP2R2B-AS1* toxicity is not entirely clear, we speculate that the toxicity may be, in part, mediated by “pure” RNA toxicity, potentially *via* the aberrant interaction between the expanded CUG repeat and RNA binding proteins (RBPs), and in support of this speculation, we observed colocalization of *expPPP2R2B-AS1* foci and MBNL1, an RBP that is involved in RNA splicing. Future *in vitro* biotinylated RNA pull down experiments^37^ may reveal additional RBPs that preferentially interact with the expanded CUG repeat on *expPPP2R2B-AS1* transcripts.

In addition, our data leads us to speculate that RAN translation of expanded polyalanine tracts from the *expPPP2R2B-AS1* transcript contributes to SCA12 neurotoxicity, as proteins or peptides containing long polyalanine tract have been previously shown to be toxic in *in vitro* cell culture or *drosophila* models^38-40^. The interrupted *PPP2R2B-AS1-Int(CUG)73* transcript is less toxic and also produces less RAN polyalanine protein, indirect evidence that RAN translation of expanded polyalanine tracts contributes to the transcript toxicity. Since all four alanine-encoding codons (GCT, GCC, GCA, and GCG) are GC-rich, it is not possible to design a long polyalanine-encoding transcript sequence without a GC-rich hairpin structure^39^, making it challenging to evaluate the relative contributions of the “pure” RNA toxicity and RAN polyalanine protein mediated toxicity. Future experiments employing additional tools, such as auxin-inducible degrons^41-43^, to specifically degrade RAN polyalanine protein, while leaving behind the *expPPP2R2B-AS1* transcript, may help answer this question.

Our qRT-PCR results indicate that exogenous *expPPP2R2B-AS1* transcripts are expressed at a lower level than *PPP2R2B-AS1* transcripts with a normal repeat, potentially due to that transcripts sequestered within CUG RNA foci may not be readily extractable, or that the presence of the expanded CUG repeat (i.e., hairpin structure) interferes with transcript expression, leading to a lower transcript level.

Our results provide further evidence that the pathogenesis of CAG/CTG repeat diseases is likely “multifactorial”, as bidirectionally expressed transcripts at the disease loci may have additive or interacting effects on pathogenesis, in addition to the potential effect of the repeat on transcription or translation. This is of relevance to the development of therapy, as it suggests that targeting sense proteins/transcripts may not always be sufficient to achieve significant therapeutic benefits. Hence, suppressing the expression of *expPPP2R2B-AS1* may offer an additional or an alternative therapeutic approach to SCA12. Given our finding that overexpression of the ZNF domains of MBNL1 decreased the expression of polyalanine RAN proteins from the *expPPP2R2B-AS1* transcripts, CUG-repeat blocking peptides based on MBNL1 ZNF domains may have a neuroprotective effect. Since SCA12 belongs to a large group of diseases that are caused by a CAG/CTG repeat expansion, the data from this study may apply to other CAG/CTG repeat diseases involving additive or synergistic mechanisms of protein and RNA neurotoxicity.

## Materials and Methods

### Brain Tissue

The high-quality human total brain RNA that was used for 3’ RACE was purchased from Clontech (Mountain View, CA). SCA12 KI-10 and KI-80 mouse models were generated using the CRISPR/Cas9 approach by replacing the mouse *PPP2R2B* exon 2 with the human *PPP2R2B* exon 7 containing either 10 or 80 CAG triplets (Li and Margolis, unpublished). SCA12 KI mice were maintained in the C57BL6/J background and bred and maintained according to the Johns Hopkins University IACUC protocol, in accordance with National Institutes of Health guidelines. Three-month-old SCA12 KI mice were deeply anesthetized, and cortices and cerebella collected. Tissues were kept at −80°C until the time of processing.

### Cell culture

Control and SCA12 patient-derived induced pluripotent stem cells (iPSCs) were generated using an episomal protocol by Cedars Sinai stem cell core, and cultured in mTeSR1 medium on Matrigel substrate. Lentiviral particles expressing NGN2 under a Tetracycline promoter were packaged in Emory Viral Vector Core, and added to iPSCs at 30:1 MOI, before puromycin (1ug/ml) was used to select iPSCs with NGN2 expression cassette integrated into the genome. Differentiation of puromycin-resistant iPSCs into NGN2 cortical neurons was as previously described^17^. SK-N-MC neuroblastoma cells (ATCC, Manassas, VA) were maintained in Dulbecco’s modified eagle medium with high glucose, supplemented with 10% fetal bovine serum and 1% penicillin/streptomycin/amphotericin B (Sigma, St. Louis, MO). Cells were transfected using TransIT-LT1 Transfection Reagent (Mirus Bio, Madison, WI) according to the manufacturer’s protocol.

### RNA extraction and Strand-specific RT-PCR (SS-RT-PCR)

Total RNAs from human iPSCs, iPSC derived NGN2 neurons, knock-in mouse brains, and transfected SK-N-MC cells were extracted by TRIZOL (Life Technologies), further purified (Monarch RNA Cleanup Kit, NEB, Ipswich, MA) and cleaned of genomic DNA (Ambion Turbo DNA-free kit, Life Technologies, Grand Island, NY). To examine *PPP2R2B-AS1* transcript expression in human cells, tissue, or mouse brains by SS-RT-PCR, 300ng or 500ng of RNA was reverse transcribed using the LK-F1 primer (SuperScript III First-Strand Synthesis System, Life Technologies), followed by a one round PCR using LK primer and a R1 primer (with manual hot start and touchdown PCR protocol) for 35 cycles. To examine *PPP2R2B* transcript expression by SS-RT-PCR, LK-R1 primer was used during the reverse transcription (RT), followed by a one round PCR using LK primer and F1 primer for 35 cycles. The PCR products were resolved on 1.5% agasrose gels. Primer sequences are listed in Table S1. Primer locations are shown in Fig. 1A.

### 3’ rapid amplification of the cDNA end (RACE)

Five ug of total RNA from human total brain (Clontech) was reverse transcribed using GeneRacer oligo-dT primer and SuperScript III First-Strand Synthesis System (Life Technologies). PCR was performed using Generacer-3’ and R2 primers with the touchdown PCR protocol, as previously described^10, 10, 13^. The PCR products were cloned into pCR4-TOPO (Life Technologies, Grand Island, NY) and sequenced.

### Real-time quantitative PCR (qPCR)

To measure gene expression levels by qPCR, 1ug of total RNA extracted from transfected SK-N-MC cells was reverse transcribed using the ImProm-II Reverse Transcription System and random hexamer primers (Promega, Madison, MI). PowerUp SYBR™ Green Master Mix (Thermo Fisher) was used for qPCR. Primers F3 and R3 were used to amplify exogenous *PPP2R2B-AS1*, while primers F4 and R4 were used to amplify exogenous *PPP2R2B-AS1-3T. GAPDH* was used as an internal control. Primer sequences are listed in Table S1. qPCR was performed using a QuantStudio 12K Flex Real-Time PCR System (Applied Biosystems, Foster City, CA).

### Fluorescent *in situ* hybridization (FISH)

The presence of foci containing expanded *PPP2R2B-AS1* in transfected SK-N-MC cells was detected using a using Cy3-labeled DNA/LNA probes (CAG)_6_-CA, which were modified at positions 2, 5, 8, 13, 16 and 19 with LNA (IDT, Coralville, IA), as previously described^25^.

### Caspase 3/7 assay

Cell viability was assessed by measuring caspase 3/7 activity (Caspase-Glo 3/7 Assay; Promega) in a 96-well format at 72 hours post-transfection as previously described^13, 26, 37^.

### DNA plasmids

An *PPP2R2B-AS1* fragment containing the region 44 bp upstream of the CTG repeat and 145 bp downstream from the repeat was PCR-amplified and cloned into pcDNA3.1(-) myc-his A vector (Life Technologies, Grand Island, NY) at EcoRV site. DNA extracted from a human SCA12 iPSC line (380i-n9) served as a template for the PCR. To obtain *PPP2R2B-AS1-Int(CTG)73* constructs, the CTG repeat region was replaced by Int(CTG)73 repeat. Int(CTG)73 fragment was synthesized by Synbio Technologies (Monmouth Junction, NJ) and cloned into pcDNA3.1(-) myc-his A vector at EcoV site. The *ATXN2-AS-(CTG)22* or *ATXN2-AS-(CTG)44* constructs were generated as previously described^13^. To examine RAN translation, *PPP2R2B-AS1-(CTG)n* inserts were PCR-amplified and cloned into the XhoI and XbaI sites of A8(*KKQEXP)-3Tf1 vector, a kind gift from Dr. Laura P. W. Ranum^44^ to produce *PPP2R2B-AS1-(CTG)n-3T* plasmids (Fig. 4A). RVG-Lamp2b-HA plasmid was obtained from addgene^45^, *PPP2R2B-Flag* and *7B7D-polySer-Myc* were generated by subcloning either full length *PPP2R2B* encoding Flag tagged Bβ1 protein, or a short *PPP2R2B* fragment containing a CAG repeat encoding a potential polySer-containing protein with a Myc tag (Zhou et al, in preparation). The GFP-MBNL1 plasmid (with pEGFP-C2 backbone) was a kind gift from Dr. Charles A. Thornton^27^. The GFP-MBNL1-Δ251-388, and GFP-MBNL1-Δ12-46 were generated as previously described^29^. The MBNL1-Δ1-240 sequence was gene-synthesized by Synbio Technology (NJ) and cloned into pEGFP-C2 plasmid at BspEI and BamHI sites to generate the GFP-MBNL1-Δ1-240 construct.

### Statistics

At least three biological replicates of each experiment were performed. Data were presented as mean ± SEM. The results were analyzed using students’ t test for comparison between two groups, or one-way analysis of variance (ANOVA) followed by Tukey post hoc test for comparison between three or more groups. Statistical significance was set at P value <0.05.

## Figure Legend

**Fig. S1.**
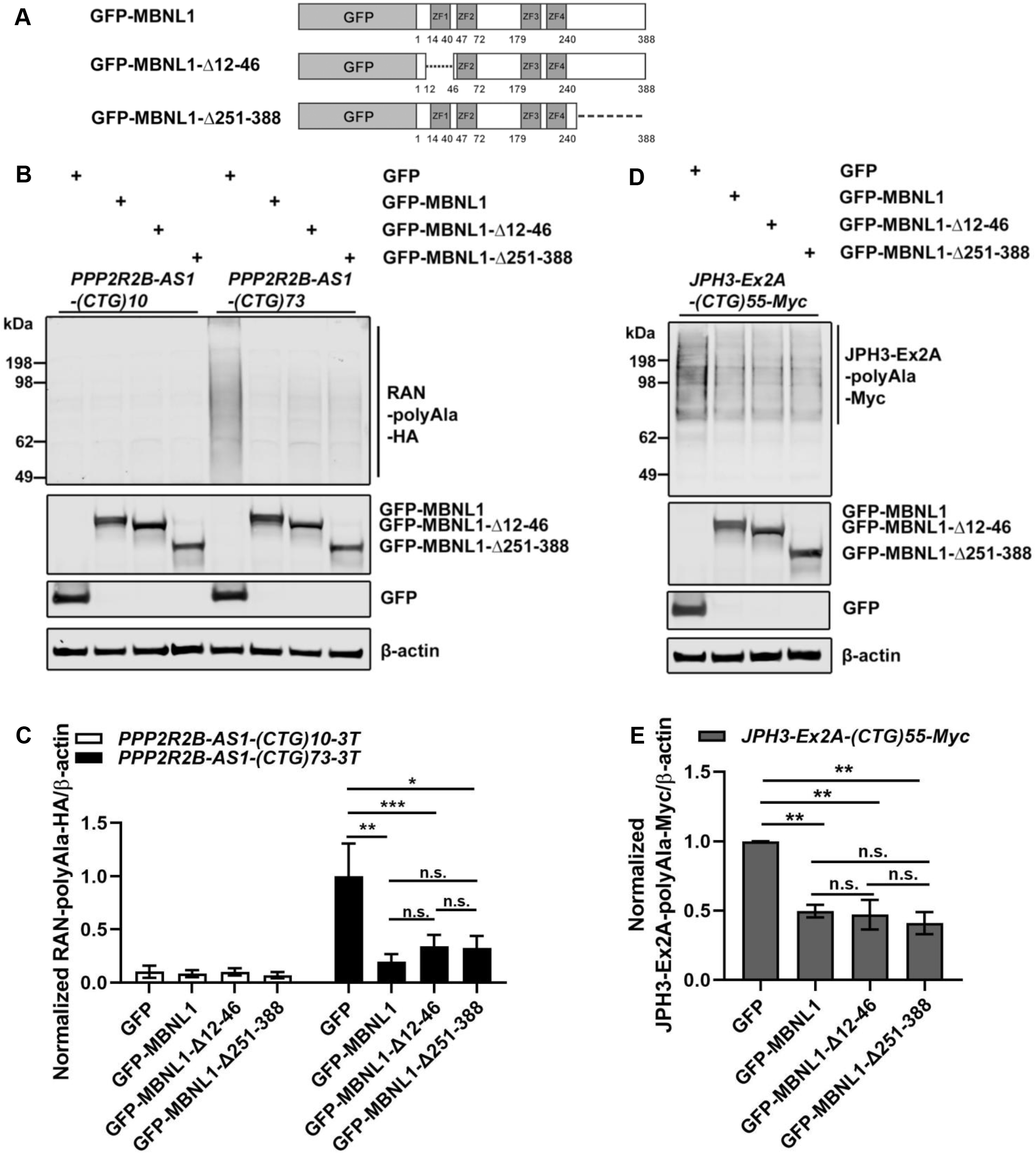
Suppression of RAN polyAla proteins from the *expPPP2R2B-AS1* transcript does not require all four ZNFs. (A). Schematic presentation of the GFP-MBNL1, GFP-MBNL1-Δ12-46, and GFP-MBNL1-Δ251-388 constructs. (B-C) Overexpression of GFP-MBNL1-Δ12-46 decreased RAN-polyAla-HA expression in *PPP2R2B-AS1-(CTG)73-3T* expressing SK-N-MC cells. N=4 biological replicates. The RAN-polyAla-HA/β-actin ratio in SK-N-MC cells co-transfected with *PPP2R2B-AS1-(CTG)73-3T* and *GFP* was normalized to 1. Mean ± SEM are shown. One-way ANOVA and Tukey posthoc test, *p<0.05, **p<0.01, ***p<0.001. (D-E) Overexpression of GFP-MBNL1-Δ12-46 decreased polyAla-Myc expression in *JPH3-Ex2A-(CTG)55-Myc* expressing SK-N-MC cells. N=4 biological replicates. The polyAla-Myc/β-actin ratio in SK-N-MC cells co-transfected with *JPH3-Ex2A-(CTG)55-Myc* and *GFP* was normalized to 1. Mean ± SEM are shown. One-way ANOVA and Tukey posthoc test, *p<0.05, **p<0.01, ***p<0.001.

**Table S1.**
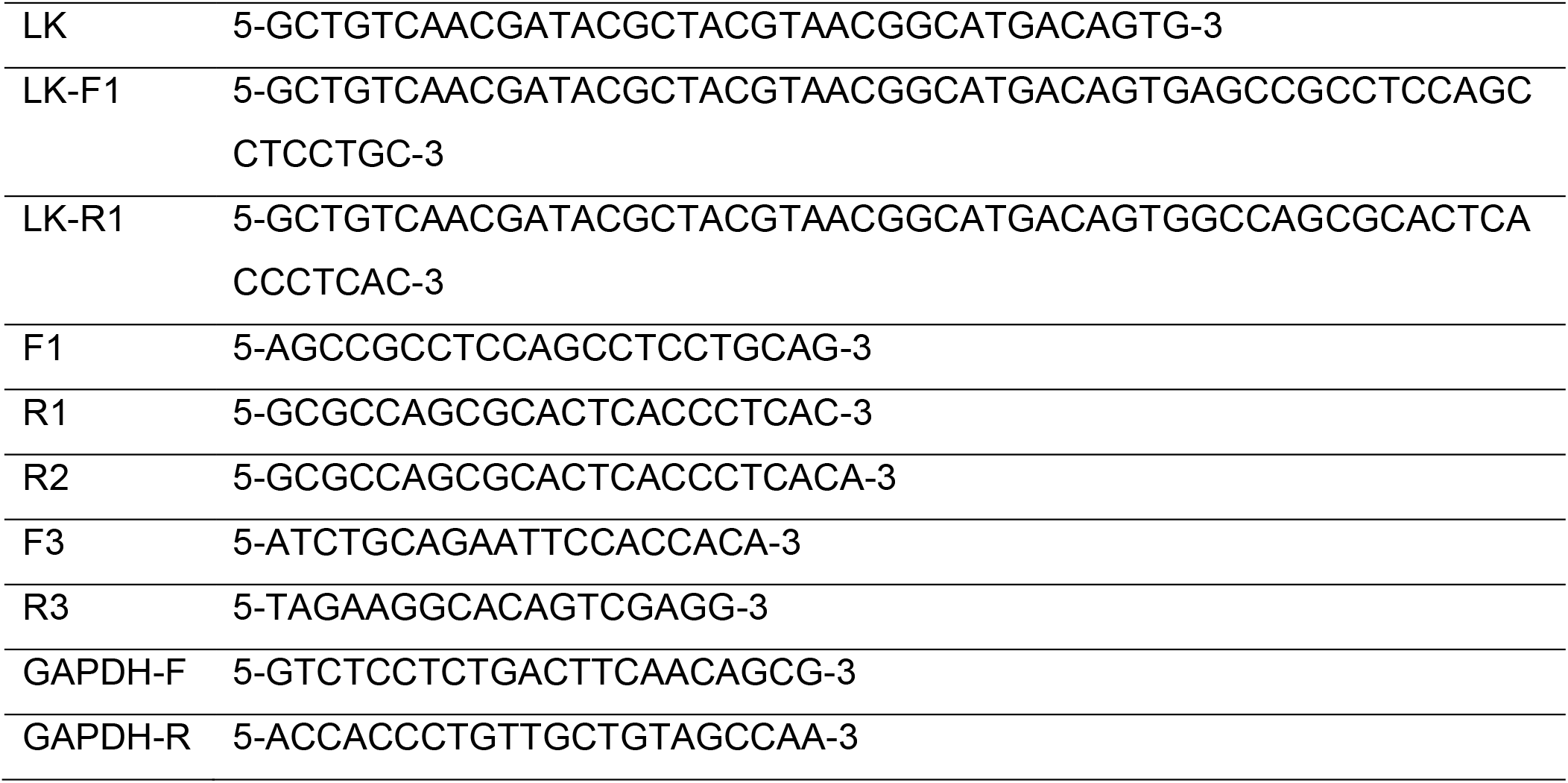
Primer sequences

## Acknowledgments

The authors thank Dr. Laura P.L. Ranum, and Dr. Charles A. Thornton for the kind gift of the A8(*KKQ_EXP_)-3Tf1, and GFP-MBNL1 constructs, respectively. This work was supported by the National Institutes of Health (NS112687, NS122756, and NS112796), the ABCD Charitable Trust, and the National Ataxia Foundation.

## Author contributions

P.P.L. conceptualized the study, and C.Z. and P.P.L. designed the experiments. C.Z., H.B.L., F.B., B.W., M.Y., R.L.M., and P.P.L. acquired and/or analyzed the data. C.Z., R.L.M. and P.P.L. wrote the manuscript. All the authors had final approval of the submitted version.

